# Unveiling *Leishmania* invasion of fibroblasts: calcium signaling, lysosome recruitment and exocytosis culminate with actin-independent invasion

**DOI:** 10.1101/420091

**Authors:** Victor Soares Cavalcante-Costa, Mariana Costa-Reginaldo, Thamires Queiroz-Oliveira, Anny Carolline Silva Oliveira, Natália Fernanda Couto, Danielle Oliveira dos Anjos, Jane Lima-Santos, Luciana de Oliveira Andrade, Maria Fátima Horta, Thiago Castro-Gomes

## Abstract

Intracellular parasites of the genus *Leishmania* are the causative agents of human leishmaniasis, a widespread emergent tropical disease. The parasite is transmitted by the bite of a hematophagous sandfly vector that inoculates motile flagellated promastigote forms into the dermis of the mammalian host. After inoculation, parasites are ultimately captured by macrophages and multiply as round-shaped amastigote forms. Macrophages seem not to be the first infected cells since parasites were observed invading neutrophils first whose leishmania-containing apoptotic bodies were latter captured by macrophages, thereby becoming infected. The fact that *Leishmania spp* are able to live and replicate inside immune phagocytic cells and that macrophages are the main cell type found infected in chronicity created the perception that *Leishmania spp* are passive players waiting to be captured by phagocytes. However, several groups have described the infection of non-phagocytic cells *in vivo* and *in vitro*. The objective of this work was to study the cellular mechanisms involved in the invasion of non-professional phagocytes by *Leishmania*. We show that promastigotes of *L.amazonensis* actively induces invasion in fibroblasts without cytoskeleton activity, thus by a mechanism that is distinct from phagocytosis. Inside fibroblasts parasites transformed in amastigotes, remained viable for at least two weeks and re-transformed in promastigotes when returned to insect vector conditions. Similarly to what was observed for *T. cruzi*, infection involves calcium signaling, recruitment and exocytosis of lysosomes involved in plasma membrane repair and lysosome-triggered endocytosis. Conditions that alter lysosomal function such as cytochalasin-D and brefeldin-A treatment or the knockout of host cell lysosomal proteins LAMP-1 and 2 dramatically affected invasion. Likewise, triggering of lysosomal exocytosis and lysosome-dependent plasma membrane repair by low doses of streptolysin-O dramatically increased parasite entry. Together our results show that *L.amazonensis* promastigotes are able to take advantage of calcium-dependent lysosomal exocytosis and lysosome-induced endocytosis to invade and persist in non-phagocytic cells.

**AUTHOR SUMMARY:** Intracellular parasites of the genus *Leishmania* are the causative agents of leishmaniasis. The disease is transmitted by the bite of a sand fly vector which inoculates the parasite into the skin of mammalian hosts, including humans. During chronic infection the parasite lives and replicates inside phagocytic cells, notably the macrophages. An interesting but overlooked finding on *Leishmania* infection is that non-phagocytic cells have also been found infected by amastigotes. Nevertheless, the mechanisms by which *Leishmania* invades non-phagocytic cells were not studied to date. Here we show that *L. amazonensis* can actively induce their own entry into fibroblasts independently of actin cytoskeleton activity, thus by a mechanism that is distinct from phagocytosis. Invasion involves subversion of host cell functions such as calcium signaling and recruitment and exocytosis of host cell lysosomes involved in plasma membrane repair and whose positioning and content interfere in invasion. Parasites were able to replicate and remained viable in fibroblasts, suggesting that cell invasion trough the mechanism demonstrated here could serve as a parasite hideout and reservoir, facilitating infection amplification and persistence.

## INTRODUCTION

The genus *Leishmania* comprises several species of intracellular parasites that cause a group of diseases collectively known as leishmaniasis. This parasitic infection is typical of tropical countries and occurs in several regions around the globe, affecting around 14 million people and generating 1 million new cases each year [1]. The disease is closely linked to poverty and is associated with malnutrition, population displacement, poor housing, immunosuppression and lack of financial resources. The outcome of the disease depends on the species and strain of the parasite and on the immunological and nutritional status of the patient. The cutaneous form of leishmaniasis is commonly caused by the species *L. braziliensis, L. major* and *L. amazonensis*, and is characterized by the formation of skin lesions that can either heal spontaneously over time or evolve to a chronic condition, which can disseminate and lead to massive tissue damage. The most severe form of the disease is known as visceral leishmaniasis, commonly caused by the species *L. donovani* and *L. infantum*, which affects internal organs such as the spleen and liver and is responsible for the majority of fatal cases.

Evolving a way to cross the host plasma membrane (PM) is a mandatory step for intracellular pathogens to establish infection. Therefore, a multitude of strategies to penetrate cells were developed by different microorganisms. Cell invasion can be accomplished through formation of a moving junction that drives parasites into cells as observed with the protozoans *Toxoplasma gondii* and *Plasmodium spp.* [2], direct injection of parasites through a specialized structure that punctures the PM as in microsporidians [3], induction of phagocytosis as in *Leishmania, Listeria, Chlamydia* and others [4] or subversion of host cell endocytic pathways as in *Trypanosoma cruzi* [5]. In the case of *Leishmania spp*., the parasite is transmitted by the bite of infected female phlebotomine hematophagous sand flies, which inject the flagellated infective promastigote forms into the mammalian host during blood meals. Once inside the mammalian host, promastigotes are ultimately captured by macrophages, which are considered to be their main host cells and in which parasites replicate as intracellular round-shaped forms, the amastigotes.

It has been reported that, before parasites reach macrophages, promastigotes are phagocytosed by neutrophils, the first immune cells to be recruited to infection site a few minutes after inoculation into the dermis [6]. Inside neutrophils, and already transformed into amastigotes, parasites are able to induce the apoptotic death of the host cell whose leishmania-containing apoptotic bodies are later captured by macrophages, which thereby become infected [7] [8]. Because in the lesions amastigotes are mainly observed inside macrophages, these cells are the most studied and the best established infection model. However, cells unable to perform classical phagocytosis, such as fibroblasts, epithelial and muscle cells, have been reported to harbor *Leishmania spp.* amastigotes *in vitro* and *in vivo* [9] [10] [11] [12]. Despite its potential importance, the mechanism by which *Leishmania spp.* invade such cells remains elusive. Therefore, we sought to investigate how the parasite invades cells unable to perform classical phagocytosis using fibroblasts and *Leishmania amazonensis* promastigotes as a model. Our results have shown that, *in vitro*, much like the related trypanosomatid protozoan *T. cruzi, L. amazonensis* subverts host cell endocytic pathways, triggering calcium signaling, lysosome recruitment and exocytosis to induce cell invasion in an actin cytoskeleton-independent fashion.

## RESULTS

### *L. amazonensis* invades mouse embryonic fibroblasts (MEFs) *in vitro*

In order to verify whether *L. amazonensis* was able to invade MEFs, the cells were incubated with *LLa*-RFP parasites for 1 h and stained with phalloidin-A488 and DAPI. Cells were analyzed by fluorescence microscopy using Zeiss-Apotome microscope to obtain confocal images. In figure 1A, a 3D reconstruction including all *z* stacks obtained for an infected cell is shown and displays the internalized parasite in the fibroblast (all stacks are provided in supplementary figure S1A). In figure 1B, a single focal plane of the same infected fibroblast shows a parasite (red) not co-localized with host cell F-actin (green), suggesting that invasion does not depend on cytoskeleton activity. Parasites were never observed co-localized with F-actin, which already suggested that cell entry does not need cytoskeleton activity - extra images of infected cells stained for F-actin are provide in Supplementary Figure S2A. To examine the kinetics of infection, we quantified infection rate by flow cytometry. Figure 1C shows that as early as 15 min about 18% of cells were RFP-positive. From 30 min to 4 h there were no substantial changes, but after 24 h, 50% of the cells were infected. Since external parasites can be easily removed by trypsin treatment, we can assume that RFP-positive cells are the infected cells.

**Fig 1.**
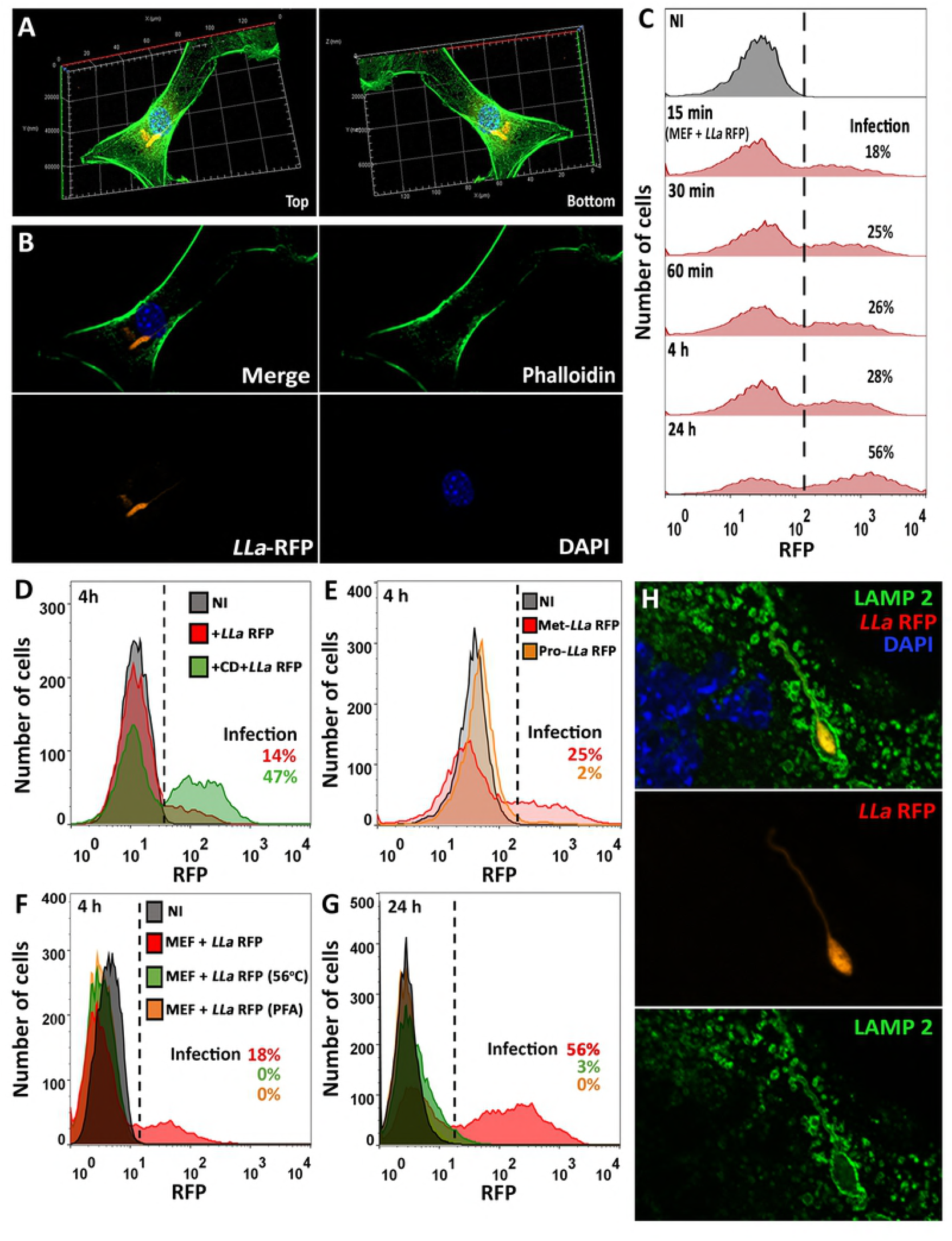
Invasion of MEFs by *L. amazonensis in vitro* depends on parasite viability and infectivity and does not require host cell actin polymerization. (A) MEF infected by *L. amazonensis*. 3D reconstruction assembled from all z-stacks obtained of an infected MEF displaying both sides of the infected cell. MEFs were incubated with *LLa*-RFP for 2 h at 37°C, labeled to visualize F-actin (green) and nuclei (blue) and imaged. (B) Single focal plane of the same infected fibroblast (Fig. 1A) shows the parasite (red) not co-localized with host cell F-actin (green). (C) Time course of MEF infection by *L. amazonensis*. Infection was performed as described in 1A, at the indicated time points cells were collected and infection quantified by FACS. Non-infected cells (NI) were gated as negative controls. (D) *L. amazonensis* infection of MEFs pre-treated with cytochalasin-D (CD). MEFs were pre-treated (green) or not (red) with 10 μM CD for 15 min, infected with *LLa*-RFP for 4 h and infection was quantified by FACS. (E) MEF infection by procyclic or metacyclic promastigotes. MEFs were infected with *LLa*-RFP metacyclic (red) or procyclic (orange) promastigotes. Infection was performed and quantified as indicated in 1D. (F and G) MEF infection with live, PFA-fixed and heat-killed *LLa-*RFP. MEFs were incubated with live (red), heat-treated (green) or PFA-fixed (orange) *LLa*-RFP for 4 (F) or 24 h (G) and infection was quantified by FACS. (H) Infected MEF with host cell lysosomal staining. Cells were labeled by immunofluorescence to visualize lysosomes (green), nuclei (blue) and imaged using Axio Imager ApoTome2 Microscope (Zeiss) to obtain a single focal plane of a MEF infected with *LLa*-RFP (red) after 2 h of infection.

To verify whether host cell actin polymerization participates in the process of invasion, MEFs were pre-treated with cytochalasin D to inhibit actin polymerization, and infection was assessed. The result (Fig. 1D) shows not only that host cell actin polymerization is dispensable for cell invasion, but that actin filament disassembly facilitates parasite entry, leading to almost four-fold increase in the infection rate. In order to determine whether invasion of MEFs is a unique property of metacyclic promastigotes, cells were incubated with either procyclic or metacyclic *LLa*-RFP promastigotes (Fig. 1E). We observed that, unlike metacyclic forms, procyclic promastigotes were not able to infect cells, indicating that the ability to invade MEFs is acquired during metacyclogenesis. To determine whether cell entry depended on the viability of parasites, MEFs were incubated with PFA-fixed or heat-treated *LLa*. We observed that, while the infection rate by living parasites reached 18% (4 h) and 56% (24 h), no PFA-fixed or heat-treated promastigotes were internalized by MEFs, apart from a negligible amount of heat-treated parasites at 24 h (Fig. 1F and G). This result showed that only living metacyclic promastigotes are able to enter MEFs.

In order to determine whether lysosomes fused with parasite-containing intracellular compartments, we stained cells with antibodies against the lysosomal protein LAMP-2 and analyzed cells by fluorescence microscopy. Figure 1H shows a single focal plane of an infected fibroblast harboring a parasite surrounded by LAMP-2 (green) after 2 h of infection, demonstrating that the parasites are fully surrounded by a membrane containing the lysosomal marker LAMP-2. Additional z stacks from this experiment are shown in supplementary figure S1B.

### *L. amazonensis* persists and replicates within LAMP-containing vacuoles inside fibroblasts

In order to evaluate the fate of the parasites internalized in fibroblasts and their ability to replicate within the host cell, we analyzed the infected population by flow cytometry after 4 and 24 h of infection. Our results showed that the RFP mean fluorescence intensity of the infected population doubled at 24 h post infection, indicating that parasites were able to replicate inside fibroblasts (Fig. 2A). To evaluate whether parasites persist inside LAMP-containing vacuoles, we performed an immunofluorescence assay in which cells infected with *LLa*-RFP were fixed, labeled with anti-LAMP-1 antibody and analyzed after 24 h of infection. Figures 2B and C show two intracellular parasites with the typical amastigote morphology inside independent LAMP2-positive vacuoles in the perinuclear region. This result shows that, upon uptake, *L. amazonensis* survives and is able to differentiate from metacyclic promastigotes into replicating amastigotes inside vacuoles with properties of lysosomes, similar to what occurs in macrophages. Images obtained by transmission electron microscopy confirmed the presence of amastigotes within host cell parasitophorous vacuoles (PV) (Figure 2D, white and black asterisks). These images revealed the characteristic subpellicular microtubules (SM) of *Leishmania* amastigotes and a close juxtaposition between the parasitophorous vacuole membrane (PVM) and parasite membranes (Fig. 2E and insert). After a week of infection amastigotes were still observed inside single PVMs (Fig. 2F), and the parasites showed no detectable alterations in their typical ultrastructural organization, such as nucleus (N), mitochondria (M) and flagellar pocket (FP) (Fig 2G). After 10 days of infection we could still observe cells containing viable amastigotes (Fig 2H and I), as demonstrated by their ability to re-transform into flagellated promastigotes (Fig. 2J) after host cells were scraped off, inoculated into promastigote culture medium and incubated at 24°C for a week.

**Fig 2.**
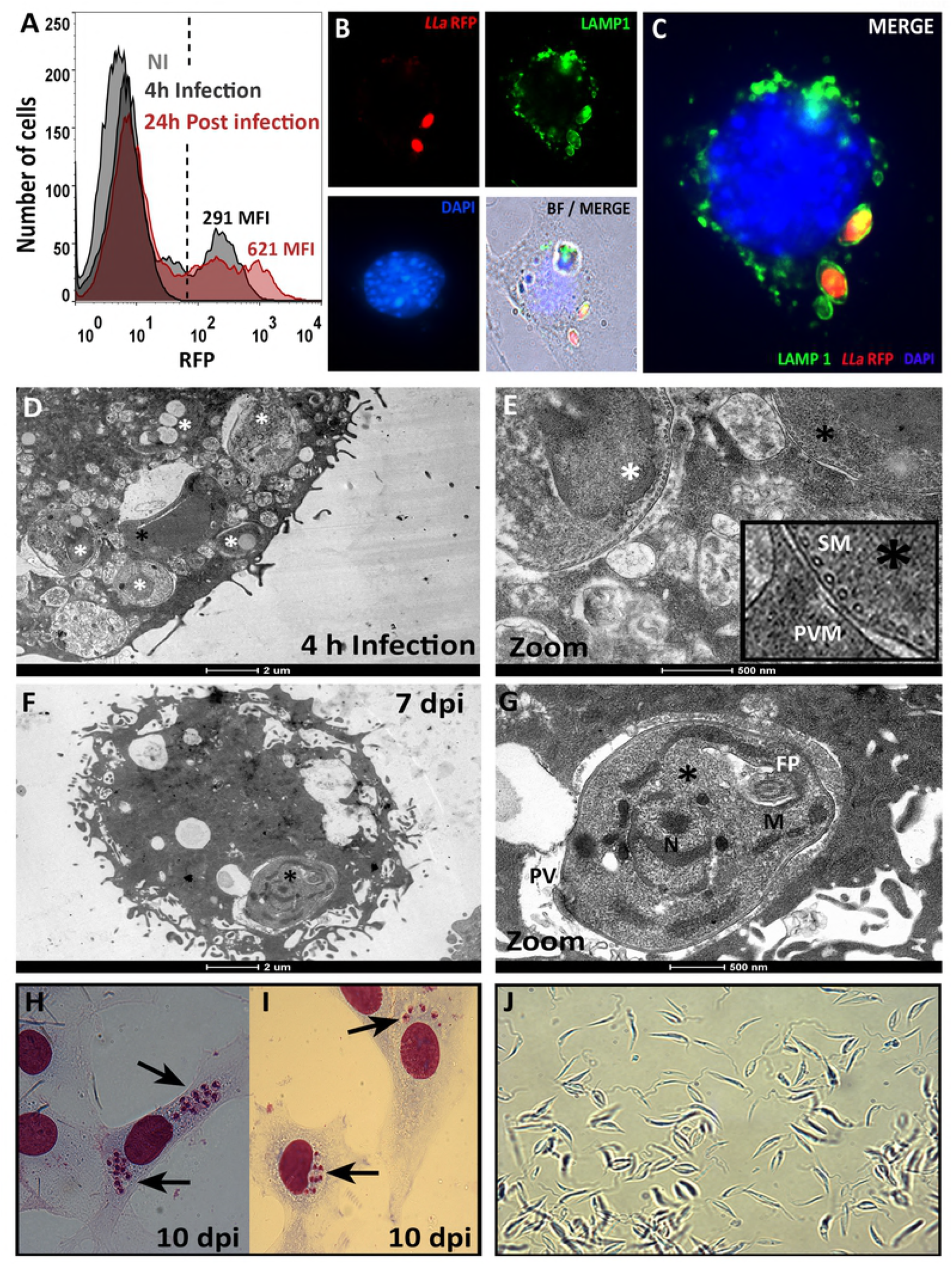
*L. amazonensis* resides in tight individual vacuoles rich in lysosomal markers and remains viable after differentiation into intracellular stages. (A) *L. amazonensis* replicates inside MEFs. After infection by *LLa*-RFP, the cell population was analyzed by FACS at 4 and 24 h post infection. The mean fluorescence intensity (MFI) of the infected population was calculated and is indicated for each curve. (B and C) *L. amazonensis* amastigotes residing in perinuclear vacuoles co-localized with lysosomal markers. MEFs were incubated with *LLa*-RFP for 24 h at 37°C and then labeled to visualize lysosomes (green) and nuclei (blue) and imaged using BX60 Upright Compound Fluorescence Microscope (Olympus). The image shows each channel individually and also merged with (B) or without the bright field (C). (D to G) Transmission electron microscopy analysis of MEFs infected with *L. amazonensis*. Cells were infected and prepared for electron microscopy after 4 h (D) or 7 days after infection (F). Asterisks show parasites inside MEFs. (E and G) Zoom-ins from the region indicated by a black asterisk in D and F, respectively. Insert in E shows a detail of the parasite PM with its typical subpellicular microtubules (SM) juxtaposed with the parasitophorous vacuole membrane (PVM). In G an amastigote is shown within the parasitophorous vacuole (PV) with the flagellar pocket (FP), nuclei (N) and mitochondrion (M). (H and I) Hematoxylin-eosin staining of MEFs 10 days after infection. (J) Promastigotes obtained from MEF-derived amastigotes. Infected MEFs shown in H and I were scraped, inoculated into insect media and imaged 10 days later by conventional light microscopy.

### *In vitro* infection of fibroblasts by *L. amazonensis* involves calcium signaling, plasma membrane permeabilization and lysosome recruitment/exocytosis

Cell invasion by intracellular parasites often involves calcium signaling, which can induce changes in the PM that promote parasite entry [13] [14] [15] [16] [5]. In order to evaluate whether *L. amazonensis* metacyclic promastigotes trigger calcium signaling in fibroblasts, we loaded MEFs with the Fluo-4AM calcium probe before inoculation of *LLa*-RFP and recorded fluorescence changes during the first 15 min of parasite-host cell contact. Intense intracellular calcium transients were detected in fibroblasts (Fig. 3A and supplementary video 1), from the first minute of incubation and continuing throughout the 15 min recorded. Figure 3B shows quantification over time of the Fluo-4AM fluorescence intensity of each indicated cell, displayed as a graphic representation of the multiple calcium transients induced in MEFs by contact with *L. amazonensis* metacyclic promastigotes. To verify whether calcium was flowing from the extracellular milieu to the cytoplasm through wounds caused by the parasites on the PM, a monolayer of MEFs was incubated with *L. amazonensis* metacyclic promastigotes in the presence of PI and then analyzed by live fluorescence microscopy. We saw that, in the presence of parasites, some host cells become PI-positive, showing that *L. amazonensis* promastigotes can induce PM permeabilization (Fig. 3C). When PI was added only at the end of the infection period and the cell population was analyzed by flow cytometry, we observed that 18% of the fibroblasts were stained by PI in the absence of calcium (Fig. 3D). On the other hand, no significant PI staining was observed when cells were exposed to the parasites in the presence of calcium (Fig. 3D), indicating that PM permeabilization is transient and that cells are able to recover when calcium is present. To evaluate whether the presence of calcium in the extracellular media is important for parasite entry, we performed the infection assay in the presence of increasing concentrations of calcium. The result (Fig.3E) shows that while in low calcium medium the infection is poor, the presence of free calcium in the media favors infection in a dose-dependent manner. Since calcium transients could also be generated intracellularly by second messengers triggered by the contact with parasites, as previously shown for *T. cruzi* and other parasites [17], the same experiment (shown in figure 3A) was performed in calcium-free medium. As observed, parasites were able to trigger calcium signaling even when calcium was absent in the extracellular media (Figs. 3 F and G). Together these results demonstrate that both intracellular calcium signaling and extracellular calcium influx occur during contact of *L. amazonensis* promastigotes and host fibroblasts.

**Fig 3.**
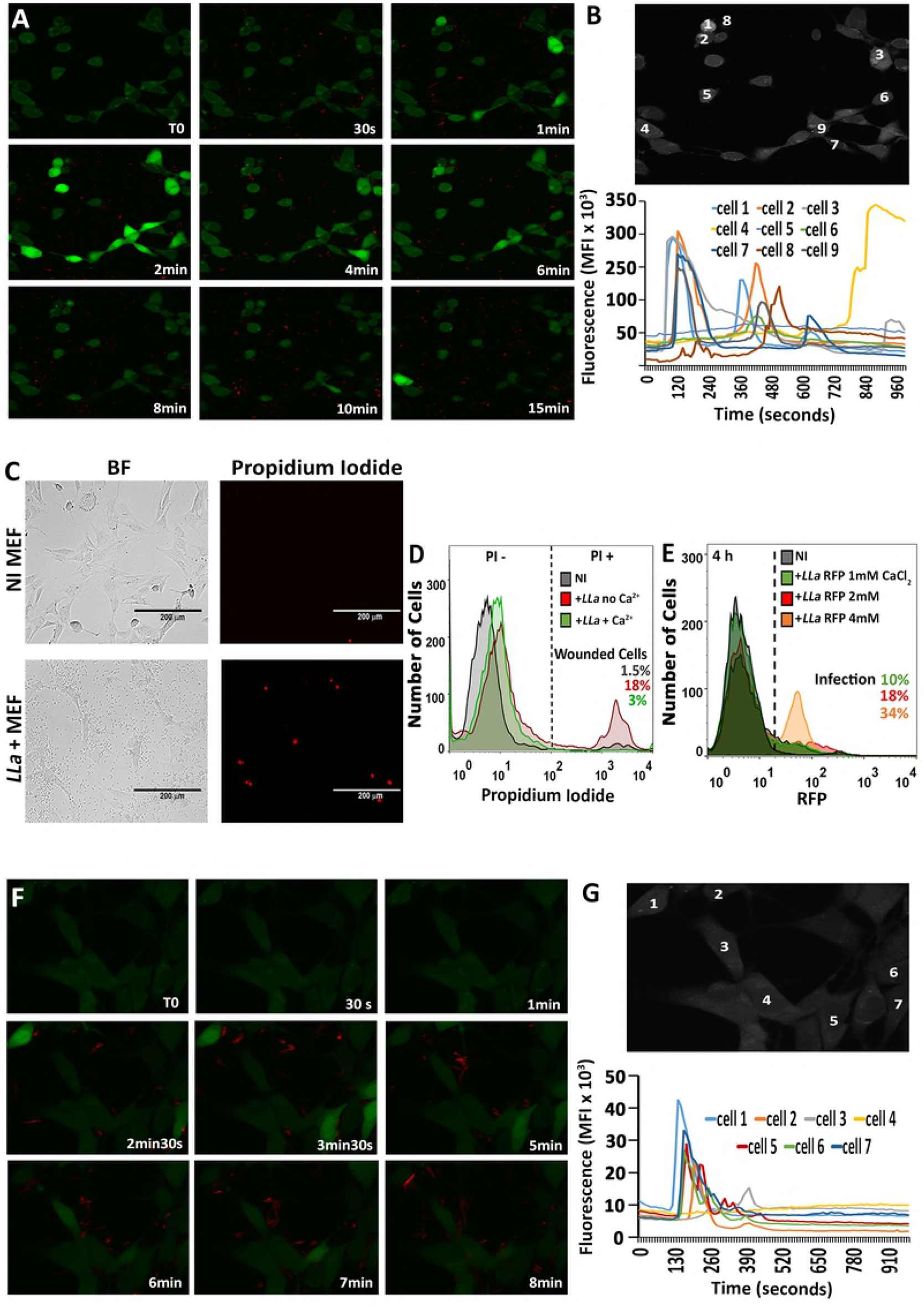
Internalization of *L. amazonensis* in MEFs involves calcium influx, PM permeabilization and intracellular calcium signaling. (A) Visualization of calcium fluxes induced by *L. amazonensis* in MEFs. MEFs were loaded with the calcium-sensitive probe Fluo-4AM and incubated with *LLa*-RFP. Cells were imaged by live confocal microscopy at 10 frames/s. (B) Graphical representation of intracellular calcium transients obtained from individual analysis of the 9 indicated cells, from the experiment in 3A. (C) Assessment of host cell PM permeability during *L. amazonensis* infection in MEFs. A MEF monolayer was incubated with *L. amazonensis* in the presence of propidium iodide (PI). After infection, the cells were examined by fluorescence microscopy. (D) Quantification of cell permeability in MEFs during *L. amazonensis* infection in the presence or absence of calcium. MEFs were incubated with *L. amazonensis* in the presence or absence of calcium for 2 h. PI was added only at the end of the experiment and the cell population was analyzed by FACS. (E) Extracellular calcium favors infection. MEFs were incubated with *LLa-*RFP in increasing concentrations of extracellular calcium for 4 h and infection was quantified by FACS. (F) Detection of parasite-induced intracellular calcium transients in MEFs. MEFs were loaded with the calcium probe Fluo-4AM and incubated with *LLa*-RFP in the absence of extracellular calcium. Cells were imaged by live confocal microscopy at 10 frames/s. G) Graphical representation of intracellular calcium transients obtained from individual analysis of 7 indicated cells, from the experiment in 3F. The videos used to build figures 3A, B, F and G are provided as supplementary data.

One of the consequences of calcium rising in the cytosol is the triggering of lysosomal exocytosis, an important step during the process of PM repair [18]. In the latter, the exocytosis of lysosomes triggers the internalization of the wounded membrane by endocytosis [19], a process that may be subverted by endoparasites to invade cells [5]. To assess whether host cell lysosomes were recruited to parasite binding site, we incubated MEFs with *L. amazonensis*, and permeabilized and labelled the cells to visualize lysosomes. The result (Fig. 4A) shows that host cell lysosomes are attracted and polarized towards parasite attachment site. To verify whether lysosomes were also exocytosing their content upon contact *L. amazonensis*, MEFs were incubated with *LLa*-RFP and then labeled with anti-LAMP-1 antibodies, this time without cell permeabilization. We observed that cells exposed luminal lysosomal protein epitopes on the extracellular leaflet of the PM (Fig.4B), indicative of lysosomal exocytosis. Quantification by flow cytometry (Fig. 4C) shows that around 30% of cells incubated with live parasites exposed LAMP-1 on their surface, an event not triggered by fixed parasites. Lysosomal exocytosis during cell entry was further confirmed by detecting beta-Hexosaminidase enzymatic activity (Fig. 4D) and the presence of acid sphingomyelinase (ASM) and cathepsin-D (Fig. 4E) in culture supernatants during host cell exposure to living *L. amazonensis* promastigotes. In order to verify whether contact with parasites also enhanced endocytosis levels in MEFs, cells were labeled with WGA-Alexa-488 to stain the plasma membrane and incubated with parasites for 15 min. After quenching the remaining extracellular fluorescence with trypan blue, the endocytosed dye was quantified by flow cytometry. The result (Fig. 4F) shows that the presence of parasites increases endocytosis in MEFs, thus making cells more susceptible to invasion.

**Fig 4.**
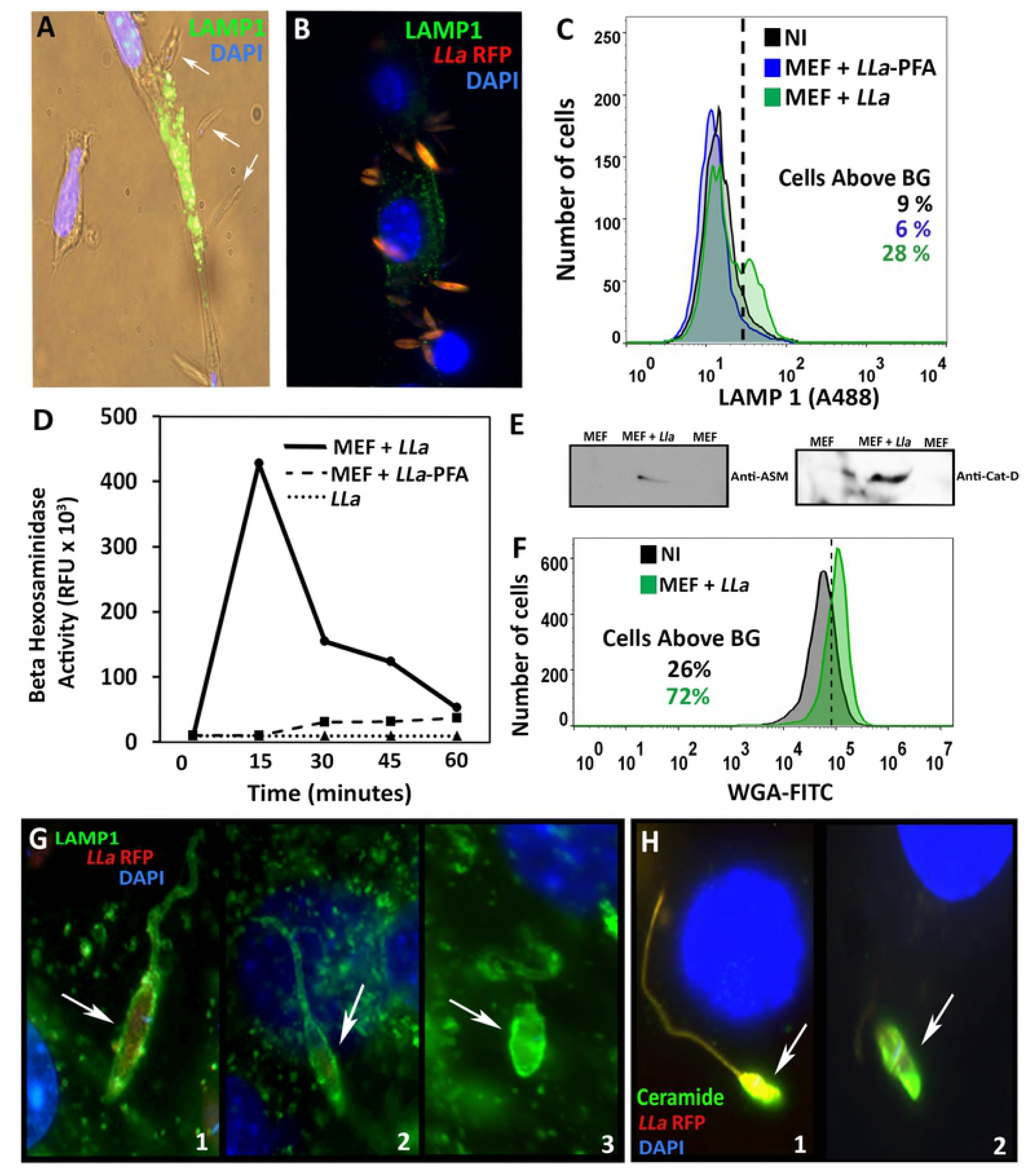
*L. amazonensis* induces lysosomal exocytosis during cell entry in MEFs. (A) Lysosomes are recruited to parasite binding site during *L. amazonensis* invasion. MEFs were incubated with *LLa*-RFP for 30 min, permeabilized and labeled to visualize lysosomes (green) and nuclei (blue), followed by fluorescence microscopy - white arrows indicate parasites interacting with the host cell (bright field) and attracting host cell lysosomes. (B) Same as A but cells were not permeabilized to visualize only the exposure of lysosomal luminal epitopes on the plasma membrane. (C) Quantification of lysosomal epitope exposure on the surface of MEFs incubated with live (blue) or PFA-fixed *L. amazonensis* (green) removed from the dish by scraping and analyzed by FACS. (D to E) Exocytosis of host cell lysosomal enzymes during *L. amazonensis* invasion in MEFs. (C) The activity of beta-Hex was assayed in supernatants of cells incubated with living (solid line) and PFA-fixed *L. amazonensis* (dashed line). Controls with *L. amazonensis* alone were carried out (dotted line). (E) Supernatants were analyzed by western blotting using anti-ASM or anti-cathepsin D (Cat-D) antibodies. (F) Endocytosis quantification in MEFs incubated with *LLa-*RFP. MEF PM was labeled with A-488-conjugated WGA before incubation with parasites for 15 min. After parasite removal the extracellular fluorescence was quenched by trypan blue and the endocytosed dye was quantified by FACS. (G to H) Detection of lysosomal markers, ceramide and LAMP recently formed *L. amazonensis* vacuoles. MEFs were infected with *LLa*-RFP for 1 h and labeled to visualize lysosomes (green) (G), ceramide (green) (H), DAPI (blue) to stain nuclei, and imaged by fluorescence microscopy.

Since exocytosis of lysosomes is followed by a massive endocytosis [20] and generates ceramide-rich vacuoles [5] in an actin polymerization-independent manner, we decided to evaluate the presence of lysosomal markers and ceramide in vacuoles of recently internalized parasites. Cells were then infected with *LLa*-RFP for 1h and labeled with anti-LAMP1 or anti-ceramide antibodies. As anticipated, parasites were completely surrounded by lysosomal markers and ceramide and both perfectly delineated bodies and flagella of the internalyzed metacyclic promastigotes (Fig. 4G and H). Conversely, and as previously stated, newly formed parasitophorous vacuoles were never covered by F-actin filaments (Fig. 1B and Supplementary Figure S2A). Together, these results indicated that the invasion process involves early lysosomal fusion and exocytosis, as previously demonstrated for *T. cruzi* [21].

### Invasion of fibroblasts by *L. amazonensis* involves the recruitment of lysosomes to the infection site to form the nascent parasitophorous vacuole

In order to follow the recruitment of lysosomes to the parasite entry site, we carried out a time-course infection of MEFs by *LLa*-RFP, and prepared cells for fluorescence microscopy using anti-LAMP-1 antibodies. At 15 min of infection, we started to observe parasites closely interacting with fibroblasts and presenting an intense co-localization with LAMP-1 at the flagellar portion (Fig. 5A). This contact with infective promastigotes creates a lysosomal polarization towards parasite binding site at the very beginning of invasion process (Fig. 4A, Fig. 5A and Supplementary Figure S2B). At 30-60 min of interaction, parasites are often observed with the flagella completely internalized and co-localized with lysosomal proteins whilst parasite body remains partially unlabeled (Figs. 5B) and surrounded by a LAMP-1-positive pocket. At 90 min, we observed parasites totally internalized, completely covered by the lysosomal marker and already located at the perinuclear region. At this point, we also started to observe the shortening of the flagella (Fig 5C). From 120 min (Fig. 5D) to 24 h (Fig. 5E), parasites were found close to the nuclei inside a juxtaposed oval-or round-shaped vacuole, completely surrounded by the lysosomal protein and with no detectable flagella, in a typical amastigote morphology.

**Fig 5.**
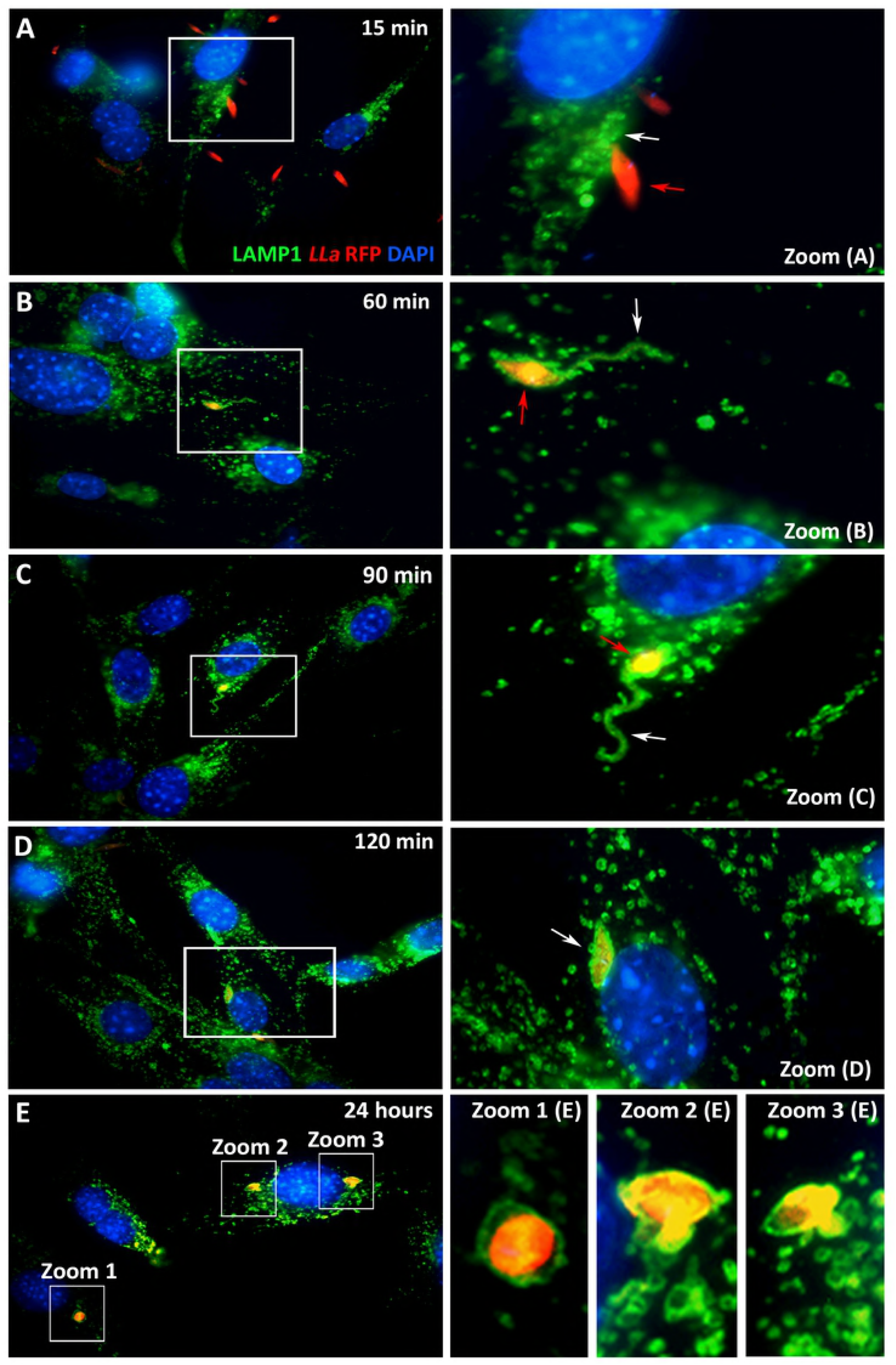
Lysosomes are recruited at early steps of *L. amazonensis* infection in MEFs and envelop parasites as they gradually transform into intracellular amastigotes. MEFs were incubated with *LLa*-RFP (red). At indicated time points, infection was stopped, cells were fixed, labeled to visualize lysosomes (green) and nuclei (blue) and imaged using BX60 Upright Compound Fluorescence Microscope (Olympus). Each figure shows the three merged channels. (A) Lysosomal recruitment to infection sites. The white arrow indicates the flagellar region and the red arrow the parasite body. (B) A partially internalized parasite with the flagella totally surrounded by the lysosomal marker LAMP-1. (C) Completely internalized parasite located at the perinuclear region displaying flagellar shortening. (D) Internalized parasite presenting an ovoid form. (E) Typical amastigote forms within LAMP-1-rich individual vacuoles at the perinuclear region.

### Lysosomal positioning and undamaged lysosomes are essential for fibroblast invasion by *L. amazonensis*

Lysosomes can be pre-linked to the PM at the cell periphery [22] [23] and associated with microtubules [24]. In order to evaluate the role of microtubule-based movement of lysosomes in fibroblast invasion by *L. amazonensis*, we treated cells with the microtubule-blocking agent nocodazole before infection. There was no difference in invasion between cells treated or not with nocodazole (Fig. 6A), suggesting that PM-associated lysosomes might be sufficient to induce invasion. Cytochalasin D and brefeldin-A are drugs known to lead to lysosome accumulation at cell periphery [21]. MEFs previously treated with each of these drugs showed a massive increase in infection (Figs. 6B and C). However, this increase was markedly blocked by nocodazole treatment (Figs. 6B and C). Cytochalasin D and brefeldin-A treatment not only led to an increase in infected cells but also to a higher number of parasites per cell, as we could observe by fluorescence microscopy (Fig. 6D, 6E and 6F) and measure by flow cytometry, which showed about 2-fold increase in mean fluorescence intensity (not shown).

**Fig 6.**
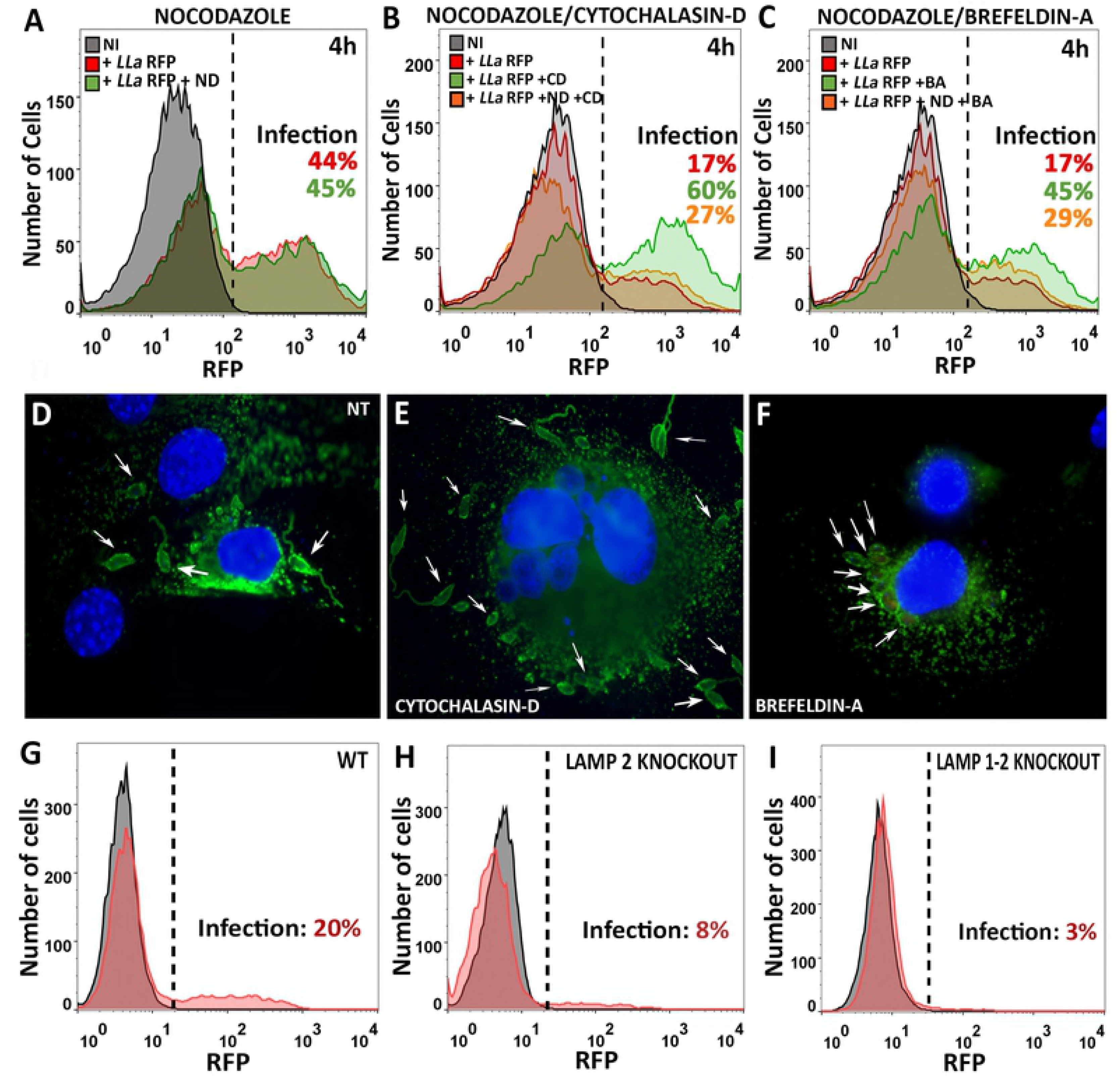
Host cell lysosome positioning and lysosomal content are crucial for MEF invasion by *L. amazonensis*. (A) Role of host cell microtubules in the invasion of MEFs by *L. amazonensis*. MEFs were pre-treated with 20 μM nocodazole for 20 min, after drug removal cells were incubated with *LLa*-RFP for 4 h at 37°C and infection was quantified by FACS. (B to C) Cytochalasin-D and brefeldin-A treatment potentiates cell invasion in a microtubule-dependent manner. (B) MEFs were treated (orange) or not (green) with 20 μM nocodazole for 20 min prior to treatment with 10 μM cytochalasin-D for 15 min. After drug removal infection was performed as described above and quantified by FACS. Infection of untreated MEFs by *LLa*-RFP is shown in red. (C) MEFs were treated (orange) or not (green) with 20 μM nocodazole for 20 min prior to treatment with 10 μM brefeldin-A for 30 min. After drug removal infection was performed as described above and quantified by FACS. Infection of untreated MEFs by *LLa*-RFP is shown in red. (D, E and F) Multi-infected cells visualized after cytochalasin-D and brefeldin-A treatments. (D) Non-treated cells, (E) cytochalasin-D pre-treated cells and (F) brefeldin-A pre-treated cells were infected as described above, fixed, labeled to visualize lysosomes (green) and nuclei (blue) and imaged using BX60 Upright Compound Fluorescence Microscope (Olympus). White arrows show internalized parasites. (G to I) Invasion of LAMP-2 and LAMP-1/2 knockout MEFs by *L. amazonensis*. (G) WT (H) LAMP-2 knockout and (I) LAMP-1/2 double knockout MEFs were infected by *LLa*-RFP as described above and infection was quantified by FACS.

Lysosomes are essential organelles whose exocytosis promotes the removal of PM lesions by endocytosis. To better evaluate the role of lysosomes in cell infection and specifically address whether PM repair is important for cell invasion, we performed the infection of LAMP-2 knockout and LAMP-1/2 double knockout MEFs with *LLa*-RFP. These cells are known to be deficient in PM repair due to the accumulation of cholesterol and caveolin in lysosomes and, for this reason, are less susceptible to the invasion of *T. cruzi* [25]. The results (Fig. 6G to I) show that the absence of these lysosomal proteins dramatically impairs *L. amazonensis* invasion.

### Generation of transient PM wounds during parasite-host cell interaction increases invasion

Lysosome recruitment to cell periphery and lysosomal exocytosis are events that can be triggered by transient PM disruption. Calcium influx through SLO pores, for example, leads to calcium-dependent exocytosis of lysosomes, which is followed by a massive compensatory endocytosis that removes the damaged membrane from cell surface [19]. Since we observed that parasites were inducing all these processes during cell entry, we decided to test whether inducing additional PM permeabilization during invasion would result in higher infection rates. First we established an ideal concentration of SLO to obtain the maximum PM damage (in the absence of calcium) with total cell recovery (in the presence of calcium) (Fig. 7A). Cells started to become permeabilized (PI-positive) at 50 ng/ml SLO, a concentration in which almost 100% of the cells were able to repair their PM (PI-negative) (blue curves). When MEFs were treated with repairable concentrations of SLO during a 15 min of incubation with *L. amazonensis*, infection of the cell population not only doubled (Fig 7B) but the number of parasites/cell also increased, as observed by around a 2-fold increase in the mean fluorescence intensity of each infected cell for both treatments (not shown). The massive increase in invasion provoked by SLO-treatment was also visualized when anti-LAMP-1-labeled infected cells were analyzed by fluorescence microscopy (Fig 7C to E). The results showed multi-infected cells (Fig 7D) in which parasites also subsequently transformed into the replicating amastigote forms (Fig. 7E).

**Fig 7.**
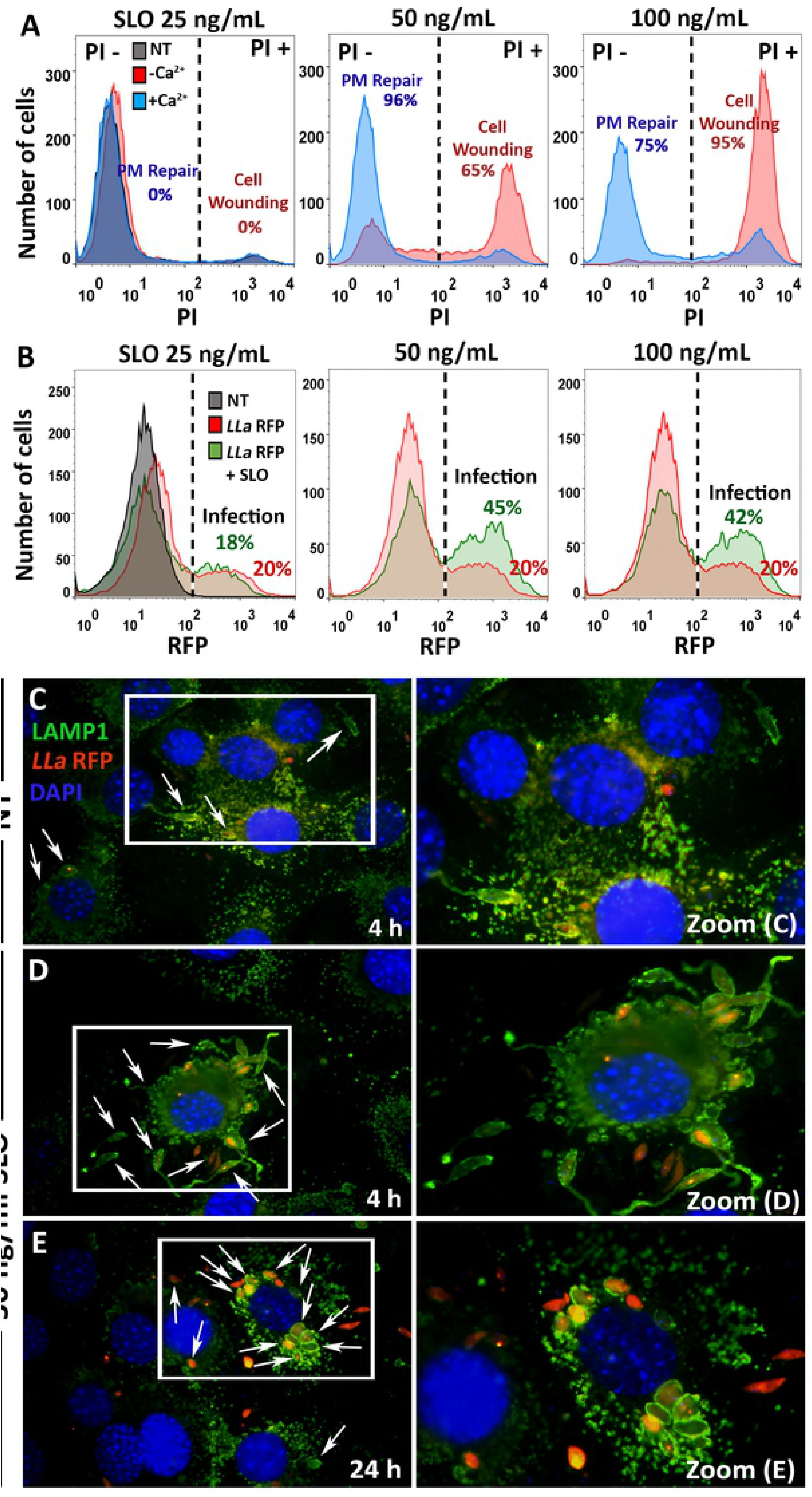
Transient PM permeabilization enhances MEF invasion by *L. amazonensis*. (A) MEFs undergo PM repair in the presence of calcium. MEFs were incubated with increasing concentrations of the pore forming protein streptolysin-O (SLO) at 37°C for 15 min in the presence or absence of calcium. After the addition of PI the cell population was analyzed by FACS. The percentages of cell wounding (red) and cells that underwent PM repair (blue) are indicated in each graph. PM repair is indicated as the percentage of cells that excluded PI after being wounded by SLO. (B) Effect of SLO-triggered PM permeabilization in the invasion of MEFs by *L. amazonensis*. MEFs were incubated with *LLa*-RFP for 4 h. At 15 min of infection SLO was inoculated in the media at the indicated concentrations and infection was quantified by FACS. Infection of non-treated (red) and SLO-treated cells (green) is shown. The percentage of infection is indicated for each curve. (C to E) MEFs multi-infected by *L. amazonensis* after SLO-treatment. The experiment showed in 7B was carried out using 50 ng/mL SLO and cells were labeled to visualize lysosomes (green) and nuclei (blue) after 4 (C and D) or 24 h (E) of infection, and imaged using BX60 Upright Compound Fluorescence Microscope (Olympus). White arrows show internalized parasites.

## DISCUSSION

The remarkable ability of *Leishmania spp.* to survive and replicate inside phagocytes, such as neutrophils, macrophages and dendritic cells, has captured most of the attention and driven nearly all research in this field during the last decades. However, these parasites are also able to infect and survive in non-phagocytic cells, a feature already observed by several authors *in vitro* and *in vivo* [9] [11] [12] [26] (reviewed by Rittig & Bogdan, 2000). In spite of the importance of such observations, almost no effort has been made to understand how these parasites succeed in infecting cells that are unable to perform classical phagocytosis. Here, using MEFs as a model, we show that entry of *L. amazonensis* into fibroblasts is a process that involves the ability of these parasites to actively induce a cell invasion mechanism involving transient PM permeabilization, calcium signaling, lysosome recruitment/exocytosis and lysosome-triggered endocytosis, much like it has been established for another trypanosomatid, *T. cruzi* [28] [21] [5]. Importantly, we demonstrated that this novel invasion mechanism by *L. amazonensis* is not a form of induced phagocytosis, since it does not seem to involve the host cell actin cytoskeleton.

While establishing assays for examining infection of MEFs by *L. amazonensis* promastigotes (Figs. 1A, 1B and 1C), it became evident that these cells could be invaded by the parasites, and that these parasites were found inside lysosome-derived vacuoles (Fig. 1H) as observed for macrophages. However, unlike the phagocytosis-mediated entry that occurs in macrophages, the invasion of MEFs by *L. amazonensis* depends on parasite direct activity, since PFA-fixed promastigotes and heat-treated parasites were not internalized (Figs. 1F and 1G). The conditions inside MEFs parasitophorous vacuoles not only allowed the typical differentiation of promastigotes into amastigotes and their replication (Fig. 2A to C), but also the persistence of viable parasites (Fig. 2H, 2I and 2J), similar to what had been described for *L. donovani* in human fibroblasts [12].

Invasion of several intracellular microorganisms such as *SalmoneLLa typhimurium* [13], group B streptococci [14], *Listeria monocytogenes* [15] and *T. cruzi* [16] [5] is accompanied by, or dependent on, a rapid increase in the levels of free intracellular calcium. In the model described here, contact with live *L. amazonensis* promastigotes also induced strong intracellular calcium transients in MEFs (Fig. 3A-B and 3F-G). Calcium seems to be an important requirement for cell invasion, since its increase in the extracellular medium positively modulated parasite entry (Fig. 3D).

We reasoned that one mechanism for the parasites to trigger calcium elevation in the cytoplasm might be the generation of host cell PM wounds during invasion. Indeed, we showed that contact with live *L. amazonensis* promastigotes wounds the PM of host cells, and the lesions are promptly repaired in the presence of calcium (Fig. 3C and 3D). In fact, when wounded, either by mechanical action or by pore-forming cytolysins, nucleated cells are able to reseal the PM in a process that involves calcium-dependent exocytosis of lysosomes [18]. Extracellularly-secreted lysosomal enzymes were proposed to act on the extracellular leaflet of the PM, triggering the removal of the wounded membrane by endocytosis [19] [29].

Calcium-dependent exocytosis of lysosomes is followed by a wave of non-conventional endocytosis [20], which is used by parasites to invade non-phagocytic cells, as previously shown for *T. cruzi* [5]. Thus, we hypothesized that host cell lysosomes are also essential for the infection of fibroblasts by *L. amazonensis*. Indeed, during infection of MEFs with *L. amazonensis*, lysosomes were recruited (Fig. 4A, Fig. 5A and Supplementary Figure 2B-15 and 30 min), fused with host cell PM (Fig. 4B-C) and exocytosed their content in the extracellular milieu (Fig. 4D-E). The exocytosis of lysosomes triggered by the parasites was followed by an increase in endocytosis levels in MEFs, indicating that the presence of parasites induces cell responses that facilitate invasion (Fig. 4F). Interestingly, recruitment of lysosomes to the infection site was observed since the very beginning of *L. amazonensis* interaction with MEFs (Fig. 4A, Fig. 5A and Supplementary Figure 2B-15 and 30 min). Notably, host cell PM wounding and exocytosis of lysosomes had already been observed even in macrophages during *Leishmania* uptake by classical phagocytosis [30], indicating that the mechanism described here might also be important during the invasion of phagocytes. However, in the case of macrophages, it was proposed that lysosomal fusion would be important to reseal PM wounds provoked by the movement of parasites after their internalization [30]. In the case presented here, exocytosis of lysosomes is an event triggered at early steps of parasite-host cell interaction and culminates with parasite internalization. Interestingly, in the experiments described here the exocytosis of the lysosomal enzyme beta-Hex peaked at 15 min of infection (Fig. 4D), matching the early triggering of calcium transients (Fig. 3A) and the appearance of infected cells as early as 15 min after parasite inoculation (Fig. 1C). It is known that after exocytosis from lysosomes, ASM cleaves sphingomyelin on cell surface producing ceramide, a lipid that promotes negative curvature of the PM enabling endocytosis [19]. A ceramide-rich vacuole, as opposed to actin-rich vacuole, is precisely what is observed in endosomes derived from the extracellular action of ASM during *T. cruzi* internalization [5]. Also similar to earlier observations, we found that recently internalized *Leishmania* parasites are surrounded by a tight PV (Fig. 2E-insert), which is intensely stained by anti-LAMP-1 (Fig. 4G) and anti-ceramide antibodies (Fig. 4H). This indicates that invasion actually takes advantage of exocytosis of lysosomes, which provide the membrane that allows parasite entry, in a mechanism that is markedly distinct from classical parasite internalization by phagocytosis in macrophages. This is corroborated by the facts that *L. amazonensis* parasites can still invade MEFs pre-treated with cytochalasin-D (Fig 1D), and that recently internalized parasites do not co-localize with actin filaments (Fig. 1B and Supplementary Figure S2A). The involvement of lysosomes in the model of invasion described here was further confirmed by the fact that cytochalasin-D and brefeldin-A, two drugs that increase infection rates for *T. cruzi* by boosting the number of peripheral lysosomes, also increased the frequency of *L. amazonensis* infection in MEFs (Fig. 6B and 6C) and the number of parasites per cell, when compared to regular infection conditions (Fig. 6E and 6F). Since both effects could be prevented by nocodazole, a drug that destabilizes microtubules and stops lysosome traffic to cell periphery [24], we can infer that microtubule-associated lysosomes may play a role in infection. Interestingly, nocodazole could not prevent infection by itself, as observed for *T. cruzi* invasion [21], which is probably due to the fact that mammalian cells already have a portion of their lysosomes pre-bound to the PM, which could be sufficient to allow parasite invasion [23]. Moreover, LAMP-2 and LAMP-1/2 knockout cells, which have modified lysosomes and impaired PM repair ability [25] were less susceptible to infection by *L. amazonensis* (Fig. 6H and 6I) when compared to wild type cells (Fig. 6G), similar to what was observed for *T. cruzi* infection with the same cell lines [25]. Additionally, our results indicate that *Leishmania* promastigotes are able to trigger calcium signaling in host cells from intracellular stores (Fig. 3F and G) since signaling also occurs in the absence of extracellular calcium. Further investigation will be needed to identify the molecules involved in this signaling. However, regardless of the origin of the calcium, from extracellular influx or intracellular reservoirs, the downstream effects important for cell invasion such as lysosomal exocytosis and its derived endocytosis would be triggered.

We still do not know how parasites induce PM injury in MEFs (Fig. 3C-D). However, at least two possibilities can be raised: 1-parasite movement against the host cell PM could generate mechanical wounds, as previously proposed for *T. cruzi* [5] and 2-the parasites might secrete cytolytic molecules leading to PM permeabilization, as proposed for *Listeria monocytogenes* [15]. Since we have described that *Leishmania spp.* produce and secrete pore-forming cytolysins [31] [32] it is possible that these molecules are responsible for permeabilizing host cells during invasion. Both possibilities would trigger calcium influx, induce lysosome exocytosis and trigger endocytosis, playing a key role in promoting parasite invasion. Indeed, when additional PM wounding was induced in MEFs by adding the pore-forming protein SLO during *L. amazonensis* invasion, the frequency of infected MEFs doubled (Fig. 7A and 7B) and multi-infected cells appeared (Fig. 7D). When wounded by SLO at the concentrations used (Fig. 7A) the host cells were able to reseal their PM, allowing the intracellular development of amastigote forms (Fig.7E).

Although several authors have already reported the presence of *Leishmania spp.* amastigotes inside non-phagocytic cells *in vivo*, it is well established that, in chronic leishmaniasis, macrophages are the main cell type found parasitized. However, it has already been shown that macrophages may not be the primary cells infected at the bite site, as neutrophils [6] and dendritic cells [33] are found to be infected by promastigotes. This demonstrates that other cells may also be important to sustain the *Leishmania* life cycle. Given that the dermis, where parasites are inoculated, is rich in non-phagocytic cells such as adipocytes, striated muscle cells, epithelial cells and fibroblasts, it is tempting to speculate that promastigotes may actively induce invasion of these cells *in vivo* through the mechanism described here.

Fibroblasts are actually interesting cells to consider during *in vivo Leishmania* infection, since they are the most abundant cells at the bite site, are major producers of chemokines that attract neutrophils and macrophages, directly interact with macrophages during wound healing and have the ability to move and spread through diapedesis [34] [35]. In addition to the ability of *Leishmania* parasites to induce cell wounding and trigger endocytic repair responses, the phlebotomine vector bite site is known to be an area of intense tissue damage, largely caused by the vector proboscis that damages the surrounding tissue to increase blood supply. Thus, at the bite site, *Leishmania* parasites probably encounter several cell types that are undergoing PM repair, a process known to involve calcium influx, lysosomal exocytosis, actin cytoskeleton rearrangements and endocytosis of wounded membranes. Besides providing a safe location to evade innate immunity, the rapid invasion of non-phagocytic cells shortly after inoculation would allow for a prompt transformation into amastigote forms, which could be later transferred to macrophages or serve as parasite reservoir. The invasion process of new macrophages during chronic infection is not yet fully understood. Amastigotes have been found to be transferred from infected to non-infected cells without cell rupture, either by phagocytosis of infected apoptotic bodies [6] or by direct cell-to-cell transfer of parasites [36]. In this context, further investigation is necessary to define whether active invasion of non-phagocytic cells is an exclusive feature of *Leishmania* promastigotes, or whether amastigotes are also able to do it. Transfer of amastigotes from an infected neutrophil to macrophages, known as the Trojan Horse strategy, was proposed to be a major mechanism allowing *in vivo* invasion of macrophages by *Leishmania spp.* [7]. In this context, it is possible that not only one, but several cell types could be Trojan Horses in *Leishmania spp*. infection, notably at the early stages. Since these parasites are able to replicate inside fibroblasts, as we report here (Fig 2A) and as described by others (reviewed by Rittig and Bogdan, 2000), it is possible that a first round of replication inside these cells could be an important step leading to infection amplification, prior to macrophage invasion.

The ability to actively induce cell invasion characterized here is a neglected feature of *Leishmania spp*, probably due to the fact that these parasites have been largely perceived as passive players taken up by phagocytosis. *In vivo* experiments depicting the very first moments of natural infection are difficult to perform and have focused mainly on neutrophils and macrophages, not covering all cell types present at the infection site. Our findings emphasize the importance of performing more accurate and strictly controlled future investigations for characterizing all cell types harboring intracellular *Leishmania* during the first moments of natural infections.

## MATERIALS AND METHODS

### 1-Parasites and host cells

The PH8 (IFLA/BR/1967/PH8) strain of *Leishmania (Leishmania) amazonensis* (*LLa*) used throughout this work was provided by Dr. Maria Norma Melo (Departamento de Parasitologia, Universidade Federal de Minas Gerais, Belo Horizonte, Brazil). Parasites were grown at 24°C in Schneider’s drosophila medium (Sigma) containing 10% heat-inactivated (hi) fetal bovine serum (FBS) (GIBCO), 100 U/ml penicillin and 100 μg/ml streptomycin (GIBCO). *L. amazonensis* expressing Red Fluorescent Protein (*LLa*-RFP) were kindly provided by Dr. David Sacks (NIH, Bethesda, USA) and cultured as described by Carneiro et al., 2018. *LLa-RFP* promastigotes were grown as described for wild type (WT) promastigotes with further addition of 50 μg/ml of geneticin G418 (Life Technologies), for selection of RFP-expressing parasites. Parasites were cultured for 4-6 days, a period in which cultures become enriched in infective metacyclic promastigotes. Metacyclic forms used in experiments were separated from procyclic forms using a Ficoll gradient, as described by Späth & Beverley, 2001.

Mouse embryonic fibroblasts (MEFs), WT, LAMP-2 knockout and LAMP-1/2 double knockout cell lines were obtained from from Dr. Paul Saftig’s laboratory (Biochemisches Institut/Christian-Albrechts-Universitat Kiel, Germany). Cells were cultured in DMEM (Gibco) containing 10% hi FBS (GIBCO) at 37°C and 5% CO_2_ atmosphere. Cultures were passaged every 48 h and plated, 24 h before experiments, on culture dishes (Sarstedt) or directly on glass coverslips, depending on the experiment. Sub-confluent cultures were used for infection experiments and were analyzed either by fluorescence microscopy or by flow cytometry. In the experiments described here we used 6 well dishes (Kasvi) and plated cells 24 h prior to experiment at 3 × 10^5^ cells per well. For immunofluorescences round coverslips were placed on the well before cell platting.

### 2-Infection experiments

Purified *LLa* metacyclic promastigotes were used throughout the experiments, unless otherwise stated. Parasites were added to dish-adherent MEFs in DMEM containing 10% hi FBS (GIBCO) which were centrifuged at 500 × *g* for 10 min at 15°C to synchronize parasite contact with cell monolayers, followed by incubation at 37°C in a 5% CO_2_ atmosphere for the indicated periods of time. All experiments were performed using a multiplicity of infection of 25 parasites per MEF. For some experiments, parasites were previously fixed in 4% PFA for 15 min, or heat-inactivated for 30 min at 56°C.

### 3-Cell labeling and western blotting

#### Immunofluorescence and fluorescent probes

MEF sub-confluent monolayers were infected with *LLa*-RFP for the indicated periods of time and fixed with 4% paraformaldehyde. Preparations were blocked/permeabilized with PBS 2% BSA 0.5% saponin and incubated with any of the following antibodies or compounds: rat anti-LAMP-1 IgG (1D4B), rat anti-LAMP-2 IgG (ABL-93) (obtained from Developmental Studies Hybridoma Bank), mouse anti-ceramide IgM (C8104-50TST) (Sigma) or alexa-488-conjugated phalloidin (Life Technologies). After washing, where appropriate, preparations were incubated for 30 min with alexa-488-conjugated equivalent secondary antibodies (Life Technologies). All preparations were stained with DAPI to visualize nuclei. Coverslips were mounted on microscope slides using anti-fading Prolong-Gold (Life Technologies) and analyzed by fluorescence microscopy. Images were acquired and analyzed using Q-Capture software or Zen Software (ZEISS), depending on the experiment, as indicated. In order to evaluate the exposure of lysosomal epitopes on the PM by flow cytometry (FACS), cells were labeled as described above but without permeabilization in order to detect only extracellular epitopes. For this purpose cells were removed from the dish with a cell scraper before analysis by FACS as described below. ***Western blotting -*** Samples were prepared with reducing sample buffer, boiled for 5 min and fractionated by SDS-PAGE on 10% acrylamide gels (BioRad). After SDS-PAGE, proteins were transferred to a nitrocellulose membrane using a wet transferring apparatus (BioRad). The membrane was blocked with 5% dry milk, followed by overnight incubation with 1:500 rabbit anti-acid sphingomyelinase (ASM) IgG (Abcam cat. # ab83354) or goat anti-cathepsin-D IgG (Santa Cruz sc-6486). After washing, membranes were incubated with the equivalent secondary antibody conjugated with horseradish peroxidase (HRP) (BioRad) at 1:10.000 in 5% dry milk for 1 h. After washing, the membrane was treated with Luminata HRP substrate (Milipore) and analyzed using a LAS-3000 imaging system (Fuji).

### 3-Quantification and visualization of infection

***FACS*** - To quantify the rate of infections we took advantage of the *LLa*-RFP described above. After infection experiments, cells were washed, treated with 0.25% trypsin (Gibco) to detach cells and non-internalized parasites and then immediately analyzed the cell population by flow cytometry using a FACSCAN II (Becton Dickinson). All analyses took into account 10.000 events (MEFs) and were performed using Flow-Jo Software. ***Light Microscopy*** - Visualization of infected cells was performed using BX60 Upright Compound Fluorescence Microscope (Olympus) after staining with a hematoxylin-eosin panoptic stain kit (RenyLab) and mounting on microscopy slides with Entellan (Merk). Images were obtained using Q-CapturePro Software. ***Fluorescence Micros*copy** – Cells labeled with fluorophore-conjugated antibodies or probes were analyzed with BX60 Upright Compound Fluorescence Microscope (Olympus) or Axio Imager ApoTome2 Microscope (Zeiss) to obtain confocal images. In order to acquire single optical sections, *z* stacks were obtained in the ApoTome mode using structured illumination microscopy technology (SIM). ***Trasmission Electron Microscopy*** *-* MEFs infected with *L. amazonensis* promastigotes were fixed in 2.5% glutaraldehyde (Sigma), 0.1 M sodium cacodylate buffer (pH 7.2) for 1 h at room temperature. Cells were then washed with 0.1 M sodium cacodylate buffer, collected with a scraper and post-fixed with a solution of 1% osmium tetroxide (O_s_O_4_) (Sigma), 0.8% potassium ferricyanide and 2.5 mM calcium chloride (CaCl_2_) for 1 h. After this second fixation step cells were washed and dehydrated in a series ascending concentration of acetone (30-100%). Finally, cells were embedded in PolyBed resin at a ratio of 1:1 (acetone/resin) for 12 h, pure resin for 14 h, and polymerized for 72 h at 60°C. Thin sections were obtained with diamond knives in an ultra-microtome (Leica UC7), collected in copper grids and stained in aqueous solutions of 6% uranyl acetate and 2% lead citrate for 30 and 5 min, respectively. Samples were observed with a Tecnai G2-20-SuperTwin FEI-200 kV Transmission Electron Microscope.

### 4-Evaluation of PM wounding and repair

The occurrence of PM wounding was evaluated by determining the degree of exclusion of the impermeant dye propidium iodide (PI), added to cell cultures at 50 μg/mL. PI-treated cells were analyzed by both fluorescence microscopy (EVOS) and flow cytometry. For fluorescence microscopy experiments using PI, MEFs were plated on 6-well culture dishes and incubated with parasites in HBBS with or without calcium in the presence of PI. To quantify PM wounding by flow cytometry PI was added as indicated, cells were detached from plates with trypsin and analyzed by FACS.

### 5-Calcium-signaling experiments

MEFs (1×10^5^ cells per well) were platted in 4-chamber glass bottom dishes and loaded with the calcium probe Fluo 4 (Invitrogen) according to Luo et al., 2011, with slight modifications. Briefly, cells were washed twice with DMEM without FBS and incubated for 50 min with Fluo 4-AM loading solution (Invitrogen). Cells were then washed once with DMEM, 3 times with calcium-free HBSS and maintained in HBSS containing or not 2 mM CaCl_2_. Calcium transients were recorded by confocal video microscopy (Nikon C2) at 10 frames per second. At 40s of imaging, 5 mM ionomycin (positive control), *LLa*-RFP or HBSS (negative control) were added to the media and the videos were recorded for up to 15 min. Image analysis and quantification of fluorescence were performed using Image-J and NIS Elements (Nikon) software.

### 5-Detection of lysosomal enzymes

MEF monolayers were incubated with *LLa* in RPMI without phenol red and supernatants were analyzed for activity of the lysosomal enzyme beta-Hexosaminidase. At the indicated time points, supernatants were collected, centrifuged to remove detached cells and beta-Hexosaminidase activity was determined as described by Rodríguez et al., 1997. Briefly, 100 μl of each supernatant were incubated with 100 μl of 2 mM substrate 4-methyl-umbellyferyl-N-acetyl-b-d-glucosaminide (SIGMA) in 6 mM citrate-phosphate buffer pH 4.5 for 15 min at 37°C. The reaction was stopped with 25 μl of 2 M Na_2_CO_3_, 1.1 mM glycine and supernatants were read in a fluorimeter at excitation/emission wavelengths of 365/450 nm, respectively. ASM and cathepsin D were detected by western blotting using anti-ASM or anti-cathepsin D antibodies, respectively, under reducing conditions and with samples prepared from FBS-free supernatants after 20 times concentration in a 10 kDa cutoff Amicon^®^ Centrifugal ultra-filter unit.

### 6-Endocytosis assay

In order to evaluate endocytosis triggered in MEFs by contact with *L. amazonensis*, 3×10^5^ MEFs were plated in a 6-well dish and the outer leaflet of the PM was labeled with 1 μg/ml Alexa Fluor 488-conjugated wheat germ agglutinin (WGA) (Life Technologies) for 1 min at 4°C. Cells were then exposed or not to *L. amazonensis* promastigotes at 37°C for 15 min followed by treatment with 0.2% trypan blue (Sigma-Aldrich) for 2 min to quench the extracellular fluorescence. After washing, the cell population was removed from the dish by trypsin treatment and analyzed by FACS to detect the remaining cell-associated fluorescence corresponding to the endocytosed dye.

### 6-Streptolysin-O (SLO) and Drug Treatments

MEF monolayers were treated with 25, 50 or 100 ng/mL of the pore-forming protein streptolysin O (SLO) during infection, or 10 μM cytochalasin-D (SIGMA) for 15 min, or 10 μM brefeldin-A for 30 min, or 20 μM nocodazole for 15 min (SIGMA). All drugs were added before infection and were removed from cells after incubation so as to not interfere with parasites viability. To evaluate plasma membrane repair triggered by SLO, fibroblasts were incubated with the indicated concentration of SLO in the absence of calcium (non-repair condition) or after restoring calcium with 2mM CaCl_2_ (repair condition) – after the addition of propidium iodide cells were analyzed by FACS.

### 7-Repeats

Each experiment in this manuscript was performed in triplicates and the results shown are representative of at least three biological replicates.

## ACKNOWLEDGMENTS AND FINANCIAL SUPPORT

We would like to thank Dr. Norma Andrews for reagent donation, advice and critical reading of this manuscript, Dr. Maria Norma Mello for proofreading this manuscript, Dr. David Sacks for kindly providing RFP-expressing parasites, Dr. Paul Saftig for kindly providing cell lines, Elimar Faria for technical support, Jacob Kames and Rodrigo Silva Reston for professional English proofreading and manuscript editing. We also would like to thank CAPI (Centro de Aquisição e Processamento de Imagens) for all support with imaging and microscopy and the Flow Cytometry Laboratory-ICB-UFMG for support with all FACS analysis.

## LEGEND TO FIGURES

**Supplementary Fig S1 –** (A and B) Series of optical sections of MEFs infected by *L. amazonensis* shown in Fig 1.A-B and Fig 1H. MEFs were incubated with *LLa*-RFP for 2 h at 37°C, labeled to visualize (A) F-actin (green) or (B) lysosomes (green) and nuclei (blue). Cells were imaged by Axio Imager ApoTome2 Microscope (Zeiss) to obtain confocal images. Z-stacks were obtained using structured illumination microscopy technology (SIM).

**Supplementary Fig S2 – Recently internalized parasites do not co-localize with F-actin, attracts lysosomes to infection site and are covered by lysosomal proteins as they enter cells.** MEFs were incubated with *LLa*-RFP (red). At indicated time points, infection was stopped, cells were fixed, labeled to visualize lysosomes (green) and nuclei (blue) and imaged using BX60 Upright Compound Fluorescence Microscope (Olympus). Each figure shows the three merged channels. (A) Three different fields showing cells containing recently internalized parasites (white arrows) and without actin coating. (B) Lysosomes are recruited to infection site and enwrap parasites during cell invasion. Parasites normally undergo transformation into amastigote forms.

**Supplementary Video 1 –** To visualize calcium fluxes induced by *L. amazonensis* MEFs were loaded with the calcium probe Fluo-4AM and incubated with *LLa*-RFP in the presence of 2 mM CaCl_2_. Cells were imaged for 15 min in a Nikon C2 confocal microscope at 10 frames/s.

**Supplementary Video 2 –** To visualize calcium transients generated intracellularly by *L. amazonensis* MEFs were loaded with the calcium probe Fluo-4AM and incubated with *LLa*-RFP in calcium-free medium. Cells were imaged for 15 min in a Nikon C2 confocal microscope at 10 frames/s.

